# Bioengineered iPSC Vessels Recapitulate Human Vascular Physiological Function and Aging Phenotypes

**DOI:** 10.1101/2025.09.02.673624

**Authors:** Shun Itai, Reina Usami, Mayuko Korekata, Jumpei Muramatsu, Mikako Katagiri, Tomoko Kasahara, Seitaro Nomura, Takaaki Abe, Hiroaki Onoe, Takafumi Toyohara

## Abstract

Vascular aging contributes to multisystem diseases and limits health span. Although various animal models have contributed to aging research, their vasculatures poorly recapitulate human physiology. Even existing tissue-engineered blood vessels fail to mimic human vascular function and pathology, hindering translational advances in vascular aging studies. Here, we present a novel human physiological vascular model fabricated via the unique molding-induced circumferential alignment of human induced pluripotent stem cell (iPSC)-derived vascular smooth muscle cells with luminally seeded endothelial cells. This architecture enabled dynamic vasodiameter changes in response to vasoactive stimuli, including hormones and intraluminal pressure. Using iPSCs from a patient with Werner syndrome, the model recapitulated aging-associated phenotypes, such as hypercontractility and increased vascular compliance, possibly due to impaired nitric oxide bioavailability. Transcriptomic and metabolomic analyses revealed age-related dysregulation consistent with vascular senescence. As a key advantage of the vasculature, spatial transcriptomic analysis demonstrated upregulation of the aging marker *CDKN1A* near the lumen and downregulation of *COL6A1* and *TPM1* throughout the vessel. Treatment with mitochonic acid 5, a mitochondria-targeted compound, significantly reversed the aging phenotypes. These findings demonstrate that our engineered vascular model recapitulates key aspects of human vascular properties and provides a platform for mechanistic studies of vascular aging and drug discovery aimed at extending health span.

## Introduction

Vascular aging plays a central role in systemic organ dysfunctions and accelerates the development of atherosclerosis, which contributes to the development of life-threatening diseases such as myocardial infarction, stroke, and lower extremity arterial diseases, often culminating in heart failure, paralysis, or limb amputation, respectively^1^. This concept aligns with the long-standing clinical observation that vascular integrity is a key determinant of biological aging, a notion first articulated in the 17th century^2^. Although pharmacological interventions targeting conventional risk factors—such as renin-angiotensin-aldosterone system inhibitors for hypertension and statins for dyslipidemia—have been widely implemented and have contributed to reductions in cardiovascular mortality, atherosclerosis remains responsible for approximately one-third of global deaths and substantially limits healthy life expectancy^3,4^. Notably, a considerable residual risk of major cardiovascular events persists despite these therapeutic advances. One major unresolved risk factor is vascular aging, but its tissue-level mechanisms remain poorly understood, and no targeted therapies are currently available^5^. A key obstacle in vascular aging research is the limited availability of human vascular tissue. Although murine models have valuable insights, they do not fully recapitulate human pathophysiology because of species-specific differences in gene expression and anatomical structure^6^.

Human cell-derived tissue-engineered blood vessels (TEBVs) may provide a promising platform for investigating the mechanisms of vascular aging and related diseases^7^. However, physiologically representative TEBVs have not yet been successfully developed. Over the past few decades, various fabrication techniques, including soft lithography, 3D printing, laser processing, electrospinning, and decellularization, have been developed to construct TEBVs^8–13^. Some of these techniques have achieved the co-culture of vascular smooth muscle cells (VSMCs) and endothelial cells (ECs) to form the multilayered structure of blood vessels. However, these models have limitations in recapitulating the function of human arteries, particularly with respect to vasoconstriction and vasodilation, core functions that regulate blood pressure and flow to maintain systemic homeostasis^14^. Two major hurdles remain in the development of physiologically relevant TEBVs. First, the circumferential alignment of VSMCs in the tunica is essential for regulating vascular tone^15^; however, previous models cannot effectively replicate this architecture. Although scaffolds with parcellated grooves promote the self-alignment of VSMCs^16–18^, these models cannot reproduce the contractile behavior of VSMCs under physiological conditions because the tissues are fixed to a rigid surface. Second, the lack of flexibility in most scaffolds restricts tissue deformation, thereby limiting functional contraction and deviating from in vivo vascular behavior. Overcoming these two limitations is critical because vasoconstriction and vasodilation are fundamental to vascular function and pathophysiology.

Here, we propose a novel rolling-up molding method of VSMCs to generate physiologically deformable TEBVs from human iPSCs, which we have named Vascular iPSC Tube Achieving Physiological deformation (ViTAP, **Figure 1(a)**). An axially aligned VSMC fiber formed with a microchannel mold is rolled up to form a helical shape with circumferentially aligned VSMCs. The rolled-up VSMCs are embedded in collagen, with ECs layered inside to establish the vascular model. The circumferential orientation and high density of VSMCs in ViTAP enable the physiological deformation of vessels in response to vasoconstrictors, vasodilators, and blood pressure. Remarkably, the fabrication process does not require any external stimulation device for cell alignment. Furthermore, the vessels can be cultured long term under pulsatile perfusion using connectors. By utilizing human iPSCs, ViTAP provides a renewable, genetically defined cell source that supports patient-specific disease modeling and future applications in regenerative medicine. These advantages position ViTAP as a robust platform for recapitulating vascular phenotypes associated with aging and diseases.

**Figure 1.**
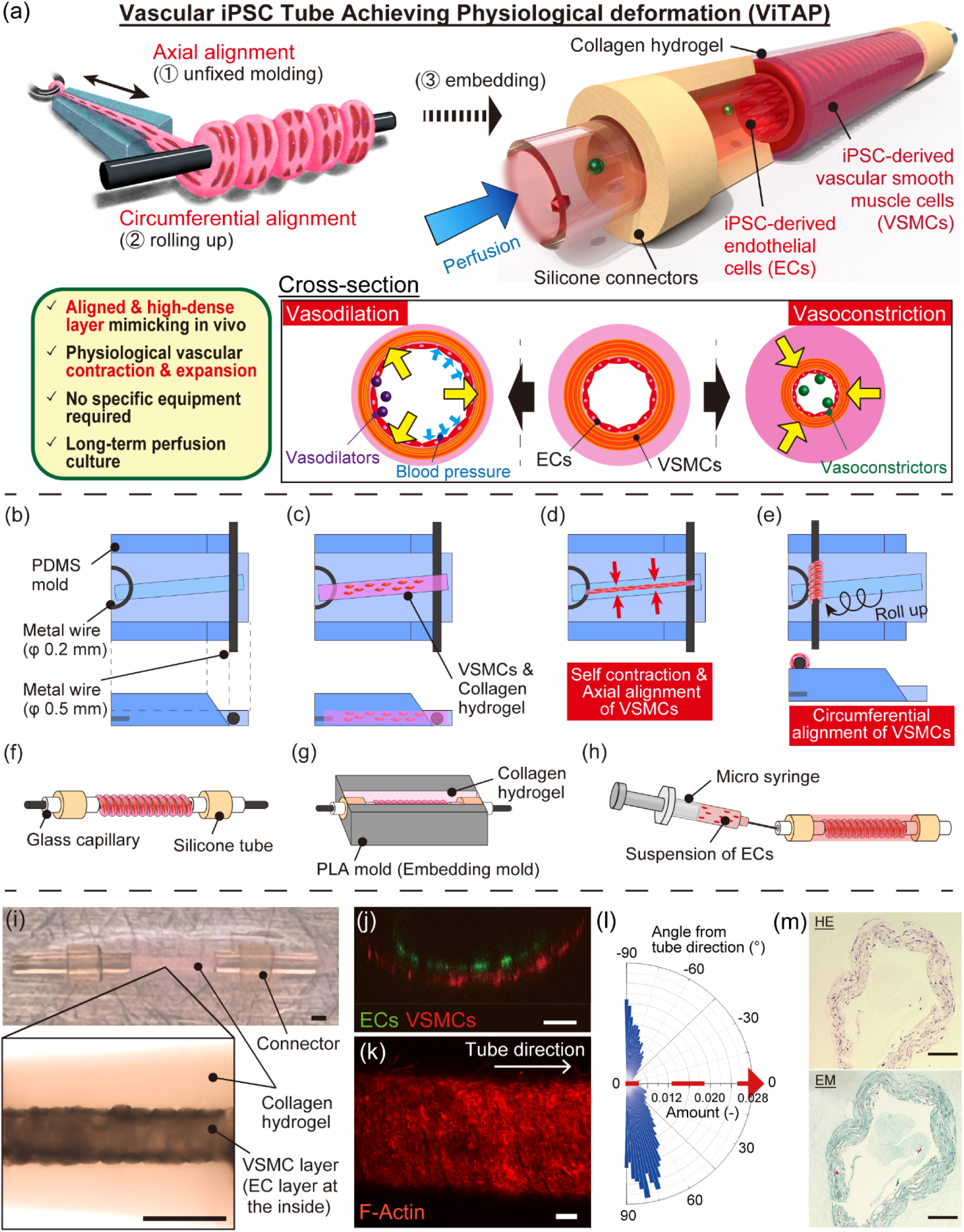
Concept of our Vascular Induced Pluripotent Stem Cell (iPSC) Tube Achieving Physiological Deformation (ViTAP) model. (a) Schematic of our model. The orientation of vascular smooth muscle cells (VSMCs), achieved through molding and rolling, facilitates physiological deformation in response to vasoconstrictors, vasodilators, and blood pressure. (b– h) Fabrication of the ViTAP model. (b) Molds were assembled. (c) VSMC-suspended collagen pre-gel solution was poured and incubated to solidify. (d) Cells were cultured for 24 h to form an aligned VSMC fiber. (e) The fiber was rolled up to form a helical shape. (f) Connectors were inserted to both sides of the tissue. (g) The tissue was embedded with collagen in a polylactic acid (PLA) mold and incubated to solidify. (h) The molds were removed, and endothelial cells (ECs) were seeded on the inner surface by injection needle to form an EC layer inside. (i) Overall and microscopic images of the fabricated model. VSMC and EC layers were embedded in collagen hydrogel, which is connected to connectors (Scale bar: 1 mm). (j) Cross-sectional fluorescent image. The co-axial layer of VSMCs and ECs was observed (Scale bar: 100 μm). (k) Fluorescent staining of actin filaments of VSMCs in the vascular tube (Scale bar: 100 μm). (l) Alignment analysis of actin filaments. (m) Hematoxylin–eosin (HE) and Elastica–Masson (EM) staining of the formalin-fixed paraffin-embedded tissue section. The high-density and multilayered structure was observed (Scale bar: 100 μm).

## Results

### Fabrication of physiologically deformable TEBVs

The ViTAP model was constructed through three primary processes: fiber formation, rolling, and embedding. Initially, metal wires were placed in a mold created via microfabrication (**Figure 1(b)**). Subsequently, a collagen solution containing VSMCs was soaked into the mold and incubated until solidified (**Figure 1(c)**). After that, the cells were cultured for 24 h to form a high-density fiber (**Figure 1(d)**). Notably, the VSMCs contracted spontaneously without external force to form the fiber, aided by the anchoring force from the metal wires. The smooth muscle fiber was then shaped into a uniform helical form, aided by the precisely designed angle of the mold (**Figure 1(e) and S1**). Next, glass and silicone connectors were inserted at both ends and embedded within a collagen solution in a microfabricated mold (**Figure 1(f,g)**). Finally, the mold and metal wires were removed, and ECs were seeded into the inner wall to build the vascular model (**Figure 1(h)**). The fabrication of the tube-shaped collagen scaffold was based on our previous report^19,20^.

VSMCs and ECs were differentiated from the healthy human iPSC line (201B7) following previously reported protocols^21,22^ (**Figure S2(a–d)**). The sparsely seeded iPSC-derived VSMCs (1.1 × 10^7^ cells/mL) formed a thin fiber shape within 24 h of culture (**Figure S3(a)**). The estimated final cell density, based on the diameter of the fiber, was approximately 2.0 × 10^8^ cells/mL (**Figure S3(b)**). Observations revealed a direct connection between the collagen gel and connectors, with the cell tissue layer visible in the collagen tube (**Figure 1(i)**). The co-axial layers of VSMCs and ECs were confirmed through fluorescent staining, as shown in a confocal micrograph (**Figure 1(j)**). In terms of alignment, the actin filaments of VSMCs were stained and analyzed using an image analysis software (**Figure 1(k)**). Compared with the vascular model without VSMC alignment as a control, our presented model exhibited a vertical alignment of VSMCs relative to the tube direction, ensuring circumferential alignment (**Figure 1(l) and S3(c)**). Additionally, histological staining revealed that the tissue had a high-density, multilayered structure similar to that of in vivo tissue (**Figure 1(m)**).

The diameter and thickness of the models used in this study were designed according to arterioles in vivo^23^ (**Figure S4(a)**); however, the size can be flexibly controlled by changing the molds. The tissue size was stable, with coefficient variations lower than 5% for every thickness and diameter (**Figure S4(b)**). Additionally, the model can be fabricated using cell lines other than iPSCs and demonstrated similar stability (**Figure S4(c–f)**). The results confirm the versatility of our method for constructing vascular models varying in size, cell source, and disease type.

### ViTAP model exhibited physiological vasoconstriction and vasodilation

As a proof of concept, the ViTAP models were physiologically examined by various vasoconstrictors, vasodilators, and blood pressure levels.

The vessel exhibited visible contraction after exposure to U46619, a thromboxane A2 receptor agonist **(Figure 2(a)**, **Supplemental Movie 1**). The contraction ratio of the vessel containing VSMC and EC layers was lower than that of the VSMC mono-culture model; however, it increased to a comparable level upon exposure to 100 μM L-NAME, an inhibitor of nitric oxide synthase (**Figure 2(b)**). Given that the nitric oxide (NO) produced by ECs relieves vasoconstriction^24^, this result ensures proper hetero-cellular interaction in our vascular model. Regarding the effect of alignment, the control model (i.e., the model with non-aligned VSMCs) showed low-amplitude, uniform contraction in both axial and radial directions, whereas the aligned model exhibited high-amplitude contraction confined to the radial direction (**Figure 2(c)**, radial: 3.97% ± 1.06% versus 16.32% ± 0.93%, *p* < 0.0001; axial mean: 3.00% ± 1.73% versus 0.27% ± 0.31%, *p* = 0.007), demonstrating the functional advantage of the aligned architecture. Moreover, several hormones aside from U46619, such as endothelin-1, adrenaline, and noradrenaline, induced vessel contraction in a concentration-dependent manner (**Figure 2(d)**). Exposure to prostacyclin (prostaglandin I₂) also induced vasodilation in a concentration-dependent manner (**Figure 2(e)**, **Supplemental Movie 2**). To further evaluate vasodilation under physiologically relevant conditions, we applied a flow of pulsatile pressure corresponding to arteriolar pressure levels^25^. Under 50 mmHg, the vessel visibly expanded (**Figure 2(f)**, **Supplemental Movie 3**) and exhibited stable, reversible cycles of contraction and dilation (**Figure 2(g)**).

**Figure 2.**
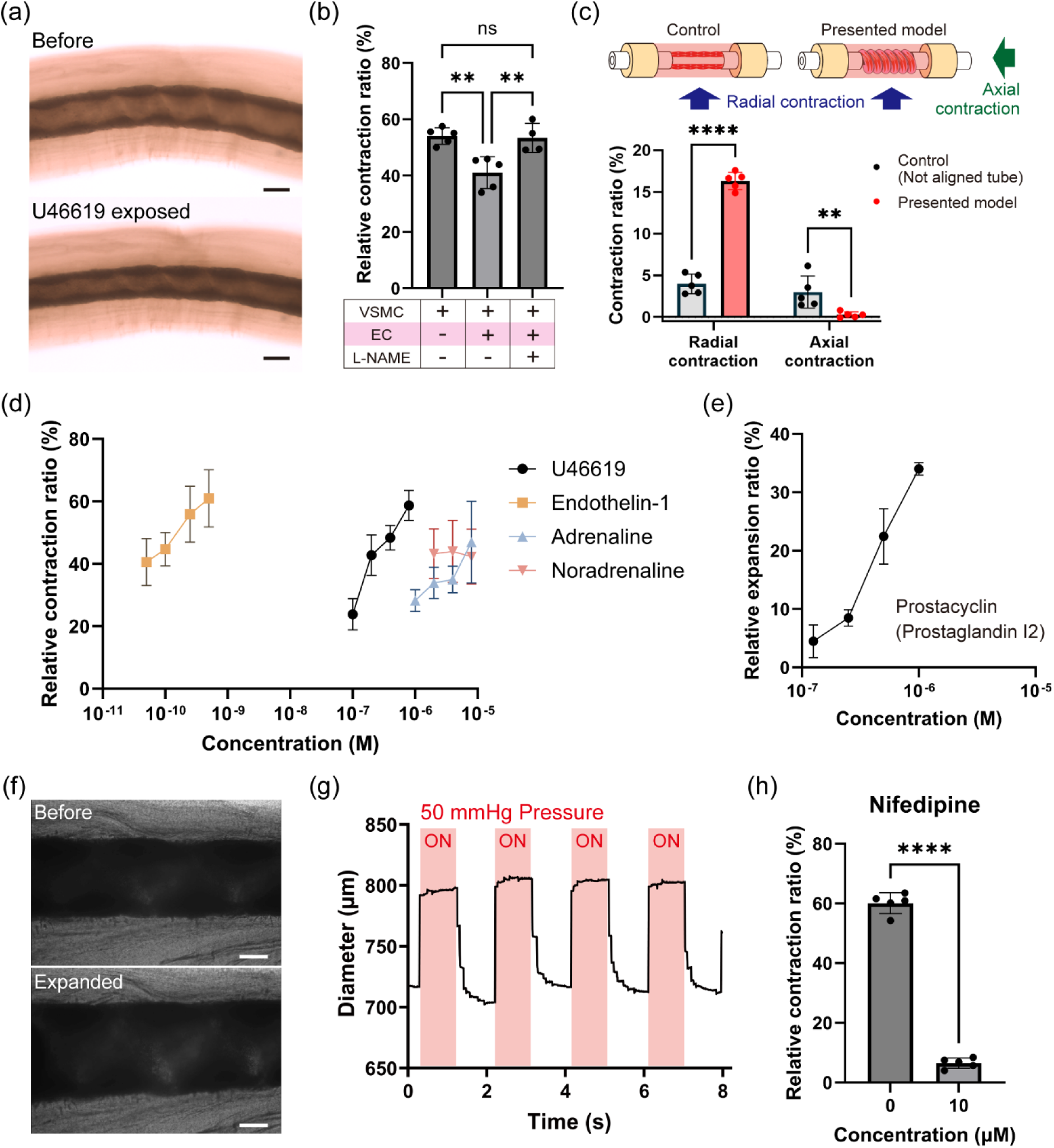
Physiological deformation of Vascular Induced Pluripotent Stem Cell (iPSC) Tube Achieving Physiological Deformation (ViTAP) model. (a) Contraction behavior in response to 200 nM U46619. The vessels showed visible contraction. (b) Relative contraction ratio of vessels to U46619 exposure normalized to 100 mM KCl (*n* = 5). Endothelial cells (ECs) alleviated the contraction by secreting nitric oxide (NO). (c) Difference in contraction behavior of vascular models by the alignment of VSMCs (*n* = 5). (d) Concentration dependency of contraction with various vasoconstrictors (*n* = 5). (e) Concentration dependency of vasodilation with prostaglandin I₂ (*n* = 5). (f) Vasodilation of the vascular model in response to 50 mmHg pulsatile blood pressure. (g) Change in the diameter of vessels to pulsatile pressure. Reversible contraction and expansion were observed. (h) Pharmacological response to nifedipine. The vasoconstriction was effectively suppressed (nifedipine: 10 μM, U46619: 200 nM, *n* = 5) (Scale bars: 500 μm, ** *p* < 0.01, **** *p* < 0.0001).

Furthermore, to evaluate pharmacological responsiveness, we pretreated the ViTAP model with 10 μM nifedipine, a calcium-channel blocker used clinically to treat high blood pressure^26^. Consistent with its vasodilatory mechanism, nifedipine significantly suppressed vasoconstrictor-induced contraction, demonstrating the functional drug reactivity of the engineered vessel (**Figure 2(h)**).

### Aged vascular model constructed from Werner syndrome iPSCs

To evaluate the capacity of the model to recapitulate senescent vasculature, we adapted the ViTAP model using WS2-C5 cells, an iPSC line derived from a patient with Werner syndrome^27^. Werner syndrome is a congenital disease characterized by premature aging induced by a mutation in the *WRN* gene, and atherosclerosis is frequently observed in patients with this disease^28^. The vascular model was successfully constructed using Werner syndrome iPSCs, which showed comparable efficiency to other cell types (**Figure S4(g–h)**). VSMCs and ECs were differentiated from iPSCs derived from a healthy individual and the patient with Werner syndrome. Vascular models were then constructed to enable phenotypic comparison (**Figure 3(a)**). Loss of *WRN* mRNA expression, consistent with the known mutation in Werner syndrome iPSCs, was confirmed by qPCR. (**Figure 3(b)**, 1.39 ± 0.26 versus 0.11 ± 0.10, *p* = 0.0119).

**Figure 3.**
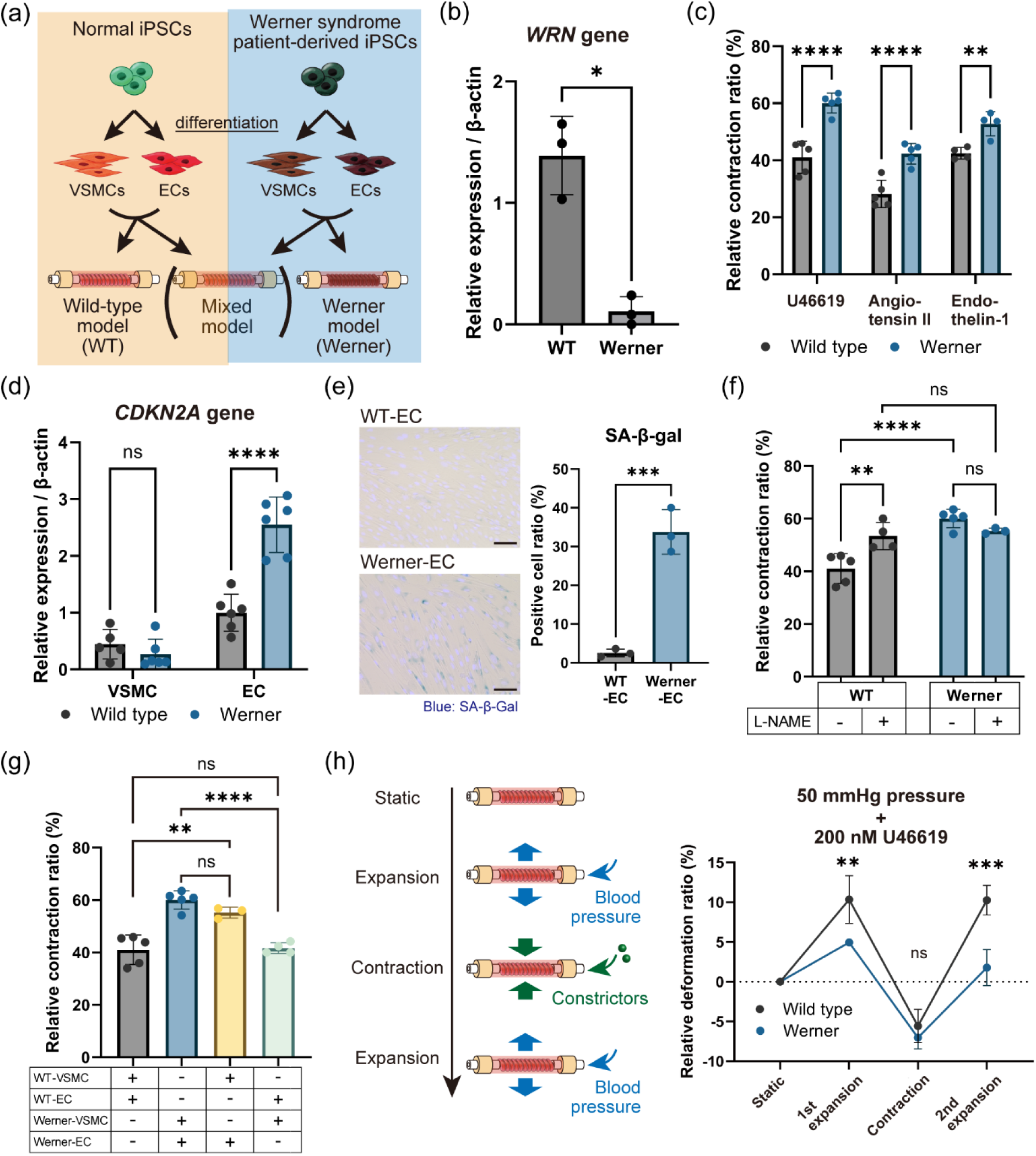
Construction of an aged vascular model and the expressed phenotypes. (a) Induced pluripotent stem cells (iPSCs) derived from a patient with Werner syndrome were utilized to fabricate the aged model. (b) qPCR results of the *WRN* mRNA expression (*n* = 3). The Werner model created with vascular smooth muscle cells (VSMCs) and endothelial cells (ECs) differentiated from Werner iPSCs exhibited proper downregulation. (c) Comparison of the contraction behavior between the wild-type (WT) and Werner models (*n* = 4 for endothelin-1, *n* = 5 for others). Excess contraction was observed in the Werner model. (d) qPCR results of the *CDKN2A* gene, an aging marker (*n* = 6). The upregulation was observed specifically in ECs. (e) Senescence evaluation by SA-β-Gal detection (*n* = 3). Werner ECs exhibited a higher positive ratio (Scale bars: 50 μm). (f) Contraction experiment under exposure of L-NAME (*n* = 4 for L-NAME sample). The increase in the contraction of the Werner model seemed to be induced by the decrease in NO production. (g) Fabrication of heterogeneous cell-source models (*n* = 4 for heterogeneous samples). (h) Comparison of the compliance. The Werner model exhibited lower compliance than the WT model (*n* = 3). (* *p* < 0.05, ** *p* < 0.01, *** *p* < 0.001, **** *p* < 0.0001)

We first evaluated the contraction responses to vasoconstrictors U46619, angiotensin II, and endothelin-1. The Werner model exhibited significantly higher contraction ratios than the healthy control model (wild type, WT) (**Figure 3(c)**, U46619: 41.06% ± 5.02% versus 60.06% ± 3.12%, *p* < 0.0001; angiotensin II: 28.19% ± 4.27% versus 42.31% ± 3.22%, *p* < 0.0001, endothelin-1: 42.45% ± 1.76% versus 52.78 ± 3.67%, *p* = 0.0061). To investigate the underlying mechanisms, we examined the senescence-associated markers in the vascular models. Notably, the expression of the *CDKN2A* gene (p16, an established aging marker^29^) was upregulated exclusively in the ECs of the Werner model, whereas VSMCs showed no significant difference in expression levels (**Figure 3(d)**, VSMC: 0.44% ± 0.23% versus 0.27% ± 0.24%, *p* = 0.6698, EC: 1.00% ± 0.30% versus 2.55% ± 0.45%, *p* < 0.0001). Senescence in the ECs derived from Werner iPSCs was further supported by positive staining for senescence-associated β-galactosidase (SA-β-gal), consistent with previous reports that ECs are susceptible to senescence^30^ (**Figure 3(e))**. Considering that one of the primary functions of ECs is to attenuate vascular contraction through NO secretion, we pretreated the models with 100 μM L-NAME prior to vasoconstrictor stimulation. Remarkably, the enhanced contraction response in the Werner model was abolished by L-NAME treatment, indicating impaired NO production as a characteristic endothelial phenotype (**Figure 3(f)**). Leveraging the modular fabrication protocol of the ViTAP model, we also constructed vascular tubes using ECs and VSMCs derived from different iPSC sources (WT and Werner). As expected, the vascular tubes containing Werner ECs showed elevated contraction ratios, irrespective of the source of VSMCs (**Figure 3(g)**).

Reduced vascular compliance is a hallmark of senescent vasculature. Increased arterial stiffness is a key feature of atherosclerosis^31^, and reduced compliance following hormone-induced contraction significantly impairs vascular functions. To assess vascular compliance, we measured vessel expansion under internal pressure simulating blood pressure before and after vasoconstriction. From a static state, the Werner model exhibited less expansion than the WT model, indicating decreased tissue flexibility. Moreover, following U46619-induced contraction, the expansion ratio declined more markedly in the Werner model than in the WT model (**Figure 3(h)**, first expansion: 10.35% ± 2.60% versus 4.96% ± 4.32%, *p* = 0.0083; contraction: −5.56% ± 1.70% versus −7.04% ± 1.15%, *p* = 0.8590; second expansion: 10.26% ± 1.52% versus 1.79% ± 1.86%, *p* = 0.0006). These findings suggest that our aged vascular model can replicate the changes in biomechanical stiffness associated with vascular aging.

### Multi-omics characterization of the aged vascular model

Transcriptomic and metabolomic analyses were conducted to assess the genetic and metabolomic phenotypes of the aged vascular model derived from a patient with Werner syndrome. Leveraging the 3D architecture of the ViTAP model, we applied spatial transcriptomics to resolve region-specific gene expression patterns within the 3D vasculatures. Our results showed that the expression of *CDKN1A* (p21, a well-established senescence marker^32^) was elevated in regions proximal to the lumen (**Figure 4(a)**), consistent with the preferential localization of senescence in ECs compared with VSMCs (Figure 3(d)). Quantitative analysis revealed significantly higher *CDKN1A* expression in EC-enriched inner regions than in VSMC-enriched outer regions in the WT and Werner syndrome vessels, supporting the spatial resolution and reproducibility of the transcriptomic data.

**Figure 4.**
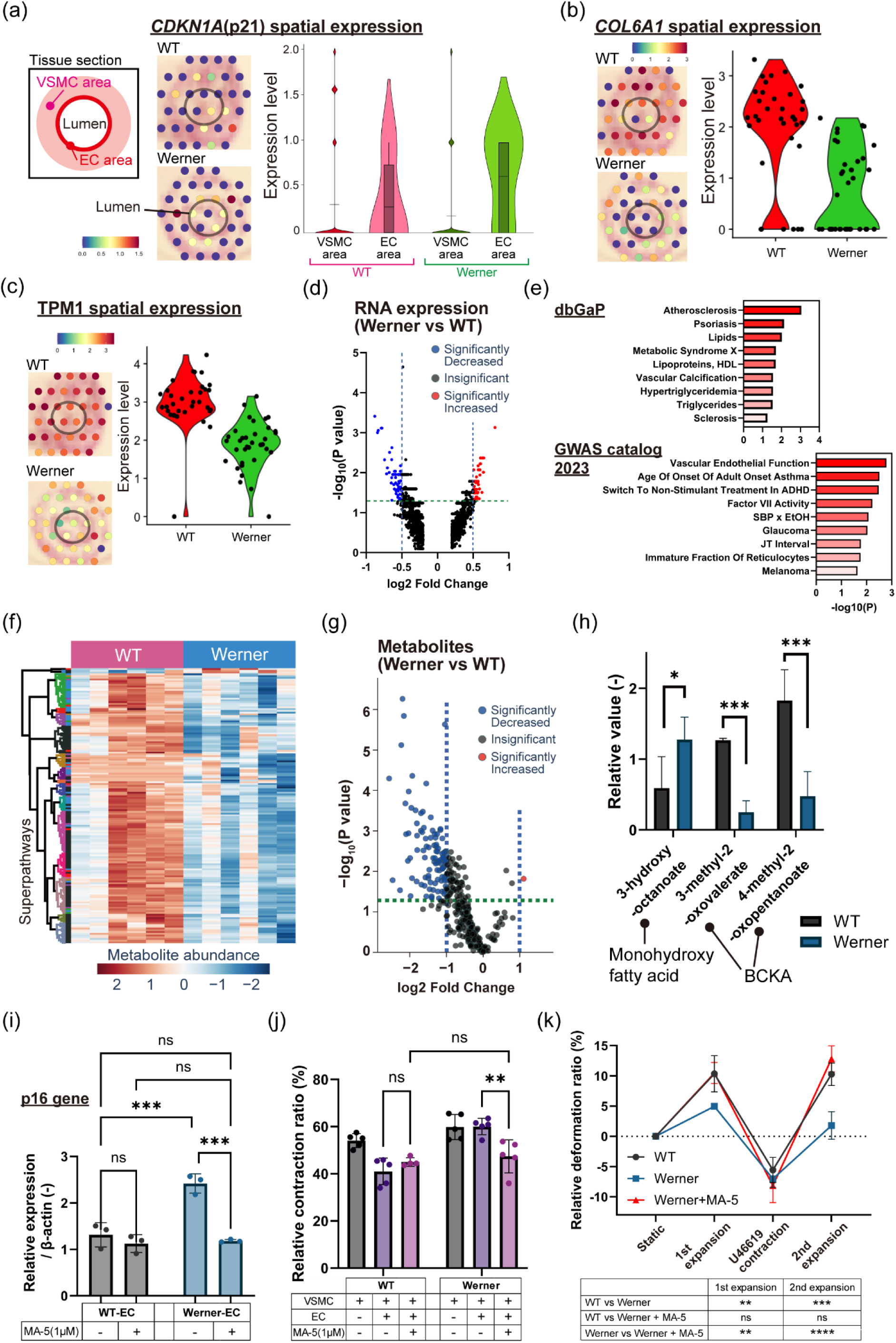
Multi-omics characterization of vascular aging in the Vascular Induced Pluripotent Stem Cell (iPSC) Tube Achieving Physiological Deformation (ViTAP) model. (a) Comparison of the spatial distribution of CDKN1A expression by spatial transcriptomics using 10X Visium. A difference was observed near the lumen area. (b) Comparison of COL6A1 expression. (c) Comparison of TPM1 expression. (d) Volcano plot comparing RNA expression levels between the wild-type (WT) and Werner models. (log_2_FC >0.2 is shown) (e) Enrichment analysis. Genes related to vascular diseases, particularly atherosclerosis, were significantly altered. (f) Metabolomic analysis of the vascular models (*n* = 6). (g) Volcano plot comparing metabolites between the WT and Werner models. Most of the metabolites were downregulated. (h) Specifically up and downregulated metabolites (*n* = 6). Metabolites in the group of monohydroxy fatty acids were upregulated, whereas those in the group of branched-chain α-ketoacids were downregulated. (i) Upregulation of the p16 gene after treatment with 1 μM MA-5 (*n* = 3). (j) Improved contraction phenotype by MA-5 (MA-5: 1 μM, U46619: 200 nM, *n* = 5). The function of endothelial cells (ECs) to secrete nitric oxide recovered. (k) Improvement of the compliance phenotype by MA-5 (*n* = 3). The treated model showed no significant difference in deformation phenotypes against the WT model. (* *p* < 0.05, ** *p* < 0.01, *** *p* < 0.001)

By contrast, several genes exhibited significantly different expression patterns across the entire vessel between the WT and Werner syndrome models. The expression levels of *COL6A1* (encoding type VI collagen) and *TPM1* (encoding tropomyosin alpha-1 chain) were significantly downregulated throughout the Werner syndrome vessel (**Figure 4(b,c)**). Type VI collagen content decreases in Werner syndrome and is associated with connective tissue abnormalities^33–35^. TPM1, which regulates VSMC contraction by facilitating actin–myosin interactions in coordination with caldesmon, has also been implicated in atherosclerosis^36–38^. Notably, although TPM1 has not been previously linked to aging, our study suggests that it plays a role in vascular aging. Downregulation of these genes may contribute to vascular aging phenotype and influence the contractile and dilatory behavior of the Werner model. These findings are consistent with prior reports suggesting that aging primarily contributes to endothelial dysfunction and EC senescence, whereas VSMCs are largely affected in terms of structural and contractile function^30^. A transcriptome-wide comparison of the entire vascular constructs further revealed multiple differentially expressed genes between the WT and Werner syndrome models (**Figure 4(d)**). Gene enrichment analysis revealed alterations in genes associated with vascular diseases, particularly atherosclerosis (**Figure 4(e)**). Furthermore, genes related to vascular endothelial function were among the top-ranked categories in a separate Gene Ontology analysis, supporting that our ViTAP-based aging vascular model recapitulates key features of vascular aging.

Comprehensive metabolomic profiling of the vascular models also revealed distinct differences between the WT and Werner models (**Figure 4(f)**). Most metabolites were significantly reduced in the aged model (**Figure 4(g)**), consistent with a previous study showing decreased metabolic activity in senescent tissues^39^. Notably, downregulated metabolites include a group of branched-chain keto acids, such as 4-methyl-2-oxopentanoate, 3-methyl-2-oxovalerate, and 3-methyl-2-oxobutyrate (**Figure 4(h)** and **S5(a)**), which are intermediates in the catabolism of branched-chain amino acids (BCAAs; e.g., leucine, isoleucine, and valine) for muscle growth and repair^40^. These findings suggest that our aged vascular model mirrors key aspects of metabolic aging, as BCAA metabolism has been implicated in aging. Conversely, metabolites belonging to monohydroxy fatty acids were upregulated in the Werner model (**Figure 4(h)** and **S5(b)**). These monohydroxy fatty acids, produced by the peroxisomal alpha-oxidation of fatty acids or branched-chain fatty acids, accumulate in senescent cells and have been associated with aging phenotypes^41^. Collectively, these results indicate that our vascular aging model can recapitulate the metabolic hallmarks of aging.

### ViTAP model demonstrated the therapeutic potential of MA-5 for vascular aging

Finally, we sought to demonstrate the utility of our Werner model as a platform for evaluating anti-vascular aging therapeutics. One promising candidate is mitochonic acid-5 (MA-5), which we previously developed and reported to enhance mitochondrial function without inducing reactive oxygen species production^42^. Although MA-5 has demonstrated anti-aging effects in *Caenorhabditis elegans* models^43^, its impact on human vascular aging has not yet been explored.

We treated iPSC-derived VSMCs and ECs with MA-5 prior to vascular model fabrication. In ECs derived from Werner syndrome iPSCs, MA-5 treatment reduced *CDKN2A* expression and restored it to WT levels (**Figure 4(i)**, WT/Werner (with MA-5): 1.12 ± 0.16 versus 1.18 ± 0.02, *p* = 0.9996). As shown in Figure 3(c), the Werner model exhibited enhanced contraction, possibly reflecting an aging-associated endothelial phenotype. Remarkably, MA-5 treatment normalized this increased contraction upon exposure to U46619 (**Figure 4(j)**, WT/Werner (with MA-5): 45.04% ± 1.57% versus 47.40% ± 6.27%, *p* = 0.9806), suggesting that MA-5 ameliorates the aging phenotype of Werner syndrome iPSC-derived 3D vasculature. While the contraction levels in the MA-5-treated Werner model remained comparable to those in the untreated models, the treated vessels demonstrated improved expansion similar to the WT model (**Figure 4(k)**). Collectively, these results indicate that our ViTAP-engineered vascular aging model is a powerful platform for evaluating anti-vascular aging therapy and underscore MA-5 as a promising novel therapeutic candidate for vascular aging.

## Discussion

To the best of our knowledge, we have developed a physiologically deformable human blood vessel model for the first time. The primary challenge in this field has been reproducing the circumferential alignment of VSMCs, an essential feature for achieving physiological vasoconstriction and vasodilation. Our ViTAP fabrication method uniquely addresses this limitation by reliably inducing circumferential alignment within a 3D construct. Recent advances in tissue engineering have produced various in vitro vascular models^8–13,44^. However, most of these models have struggled to achieve precise cellular alignment, and the scaffold flexibility needs to replicate dynamic vascular function. Another significant limitation of existing models is their inability to achieve sufficient VSMC density to generate robust contractile force because of challenges in nutrient diffusion and cell viability. Our molding and rolling method leverages the self-contraction force of cells, similar to tissue organoid formation, to create densely packed, highly aligned constructs without requiring external stimulation, such as electrical fields or mechanical stretching. This approach results in tissue architecture and functional capacity that more closely mimic in vivo vasculature than previous in vitro models. The other distinctive feature of our approach is the ability to construct vasculature using different cell sources for each layer. In the contraction experiments of the aged model, we altered the derivation of iPSCs between VSMCs and ECs within the same tissue construct, enabling the detailed dissection of the specific contributions of each cell type to the observed phenotypes. These results indicate that our vascular model functions not only as a faithful mimic of in vivo tissue but also as a higher-order system that can elucidate the intricate pathological mechanisms and identify potential therapeutic targets.

Our system offers a unique platform for comprehensively modeling disease phenotypes, providing critical insights into vascular aging and potential therapeutic strategies. Aging significantly increases the risks of numerous common diseases, including CVD, diabetes, Alzheimer’s disease, Parkinson’s disease, chronic obstructive pulmonary disease, osteoporosis, and even osteoarthritis^45^. Despite many advances in human aging research, many contributing factors remain poorly understood. Previous studies have relied primarily on models such as *C. elegans*, *Drosophila,* mice, and rabbits; thus, validation of anti-aging interventions in human-relevant models or long-term clinical studies is warranted. To obtain deeper fundamental and mechanistic insights, human in vitro models of aging must be developed^45^. In this context, our study represents the first successful development of human vascular aging in vitro model, offering a promising platform for elucidating mechanisms and developing novel therapies for vascular aging. Utilizing our 3D vascular aging model, we have recapitulated functional aspects of vascular aging and identified downregulated genes potentially involved in this process, specifically *COL6A1* and *TPM1*. The reduced expression of these genes in smooth muscle cells appears to significantly compromise tissue compliance by altering muscle fibers and extracellular matrices^33–38^. Furthermore, considering the well-established role of mitochondrial dysfunction in accelerating cellular senescence^45^, we focused on MA-5, a mitochondria-targeted therapeutic candidate. MA-5 treatment effectively ameliorated aging phenotypes in our vascular model, underscoring its potential as a novel anti-vascular aging intervention.

Our ViTAP model offers remarkable potential for vascular research and broader applications. In this study, we perfused only culture medium through the vascular construct; however, vascular aging-associated atherosclerosis in vivo involves complex interactions with monocytes and macrophages^1^. By incorporating these immune cells into the perfusion system, our model could replicate these pathological features in greater detail. Beyond blood vessels, other luminal tissues, such as the intestine, ureter, and esophagus, contain contractile muscle layers regulated by neuronal input^46^. The muscle layer must be innervated to adapt our ViTAP model for these organ systems and accurately recapitulate physiological behaviors. The model includes an ECM layer surrounding the vasculature, suggesting that neural co-culture can be achieved. Moreover, functional innervation of the muscle layer by neurons may be achieved by modulating culture conditions and cell arrangements. Incorporating neural elements into the ViTAP model would enable not only the fabrication of diverse organ constructs but also the modeling of various diseases. Leveraging these advances in tissue engineering and iPSC technologies, our ViTAP model holds significant promise for advancing detailed biomedical research and facilitating the development of new therapeutic strategies.

## STAR Methods

### Key resources table

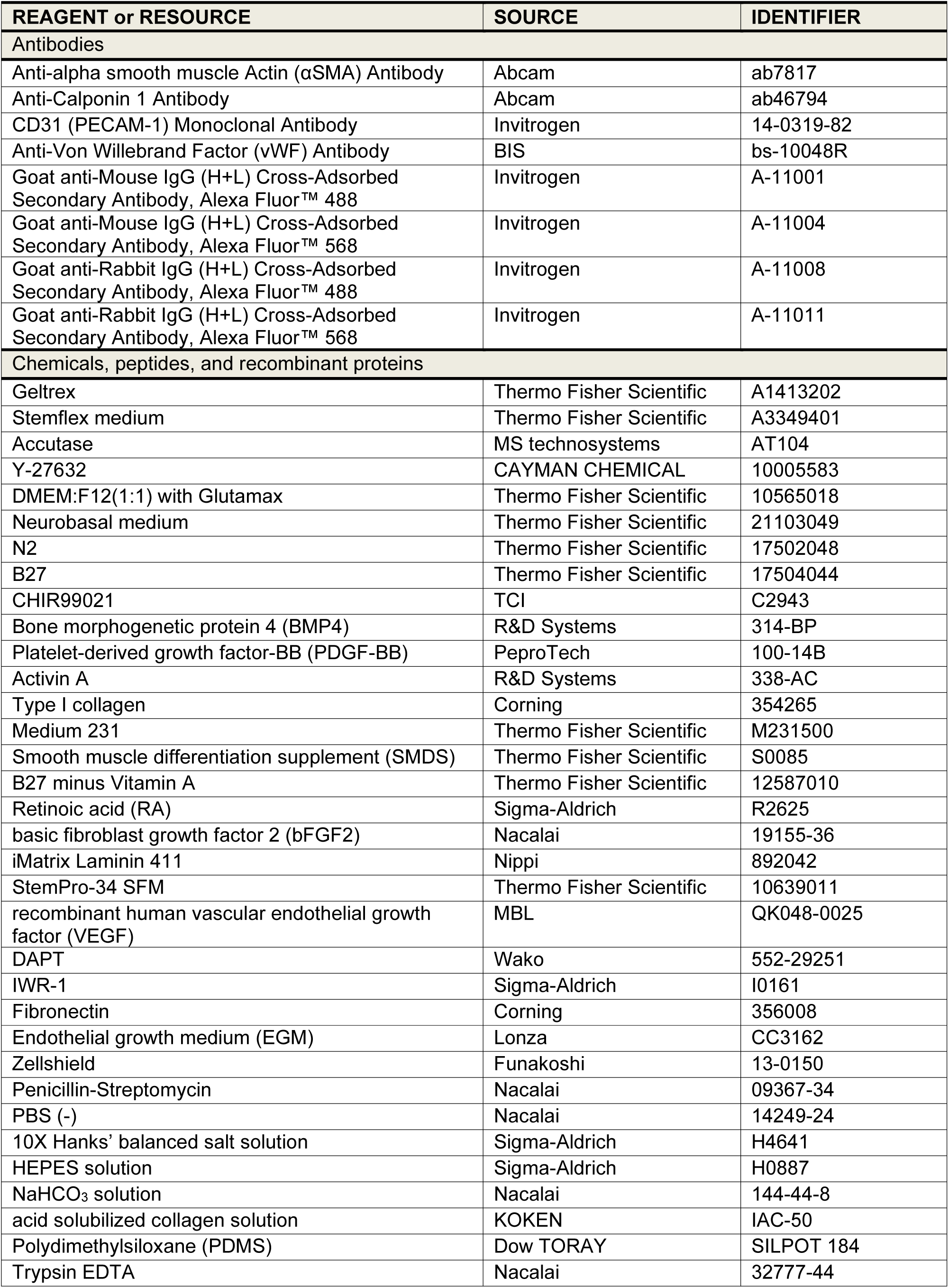
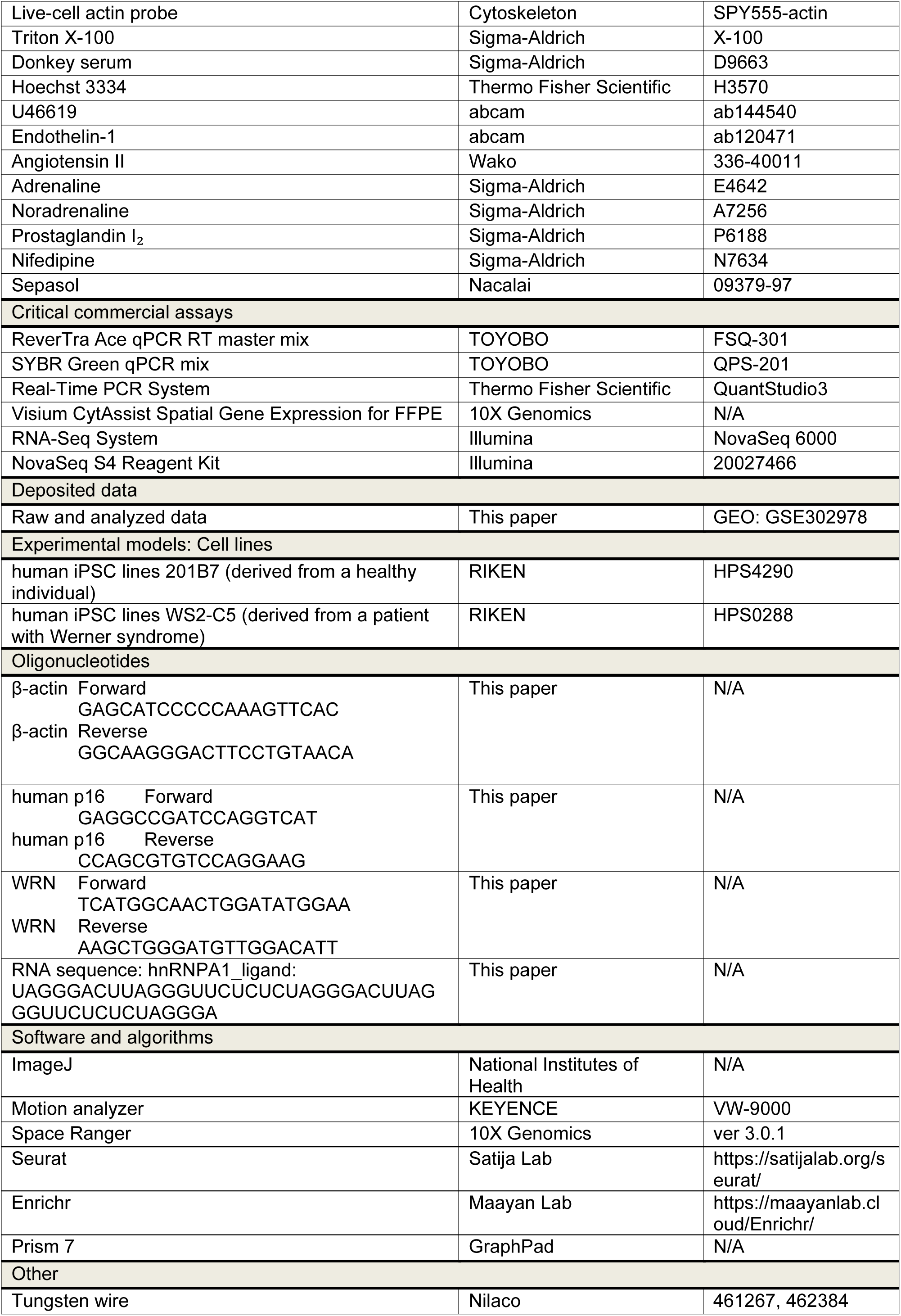
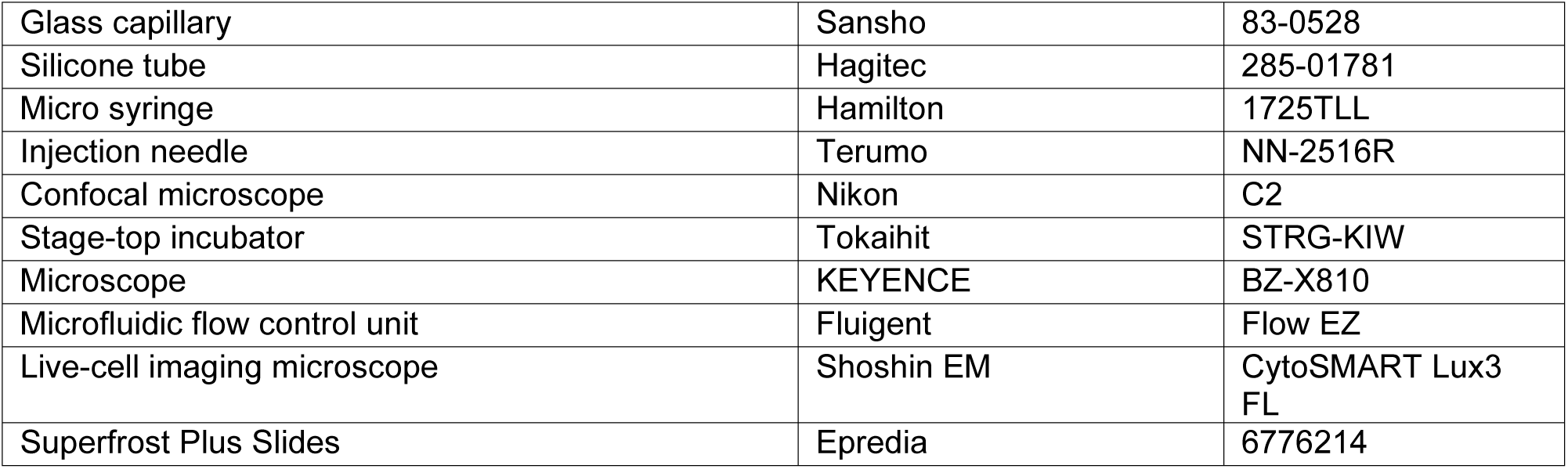

### Cell culture and differentiation of iPSCs to VSMCs and ECs

The human iPSC lines 201B7 (derived from a healthy individual, HPS4290) and WS2-C5 (derived from a patient with Werner syndrome, HPS0288) were obtained from RIKEN CELL BANK, Japan. Both human iPSC lines were maintained on Geltrex (A1413202; Thermo Fisher Scientific, MA, USA)-coated dishes in Stemflex medium (A3349401; Thermo Fisher Scientific). Cells were replated using Accutase (AT104; MS Technosystems, Japan) and seeded at a density of 10,000–20,000 cells/cm^2^ for passage.

For differentiation to VSMCs, iPSCs were dissociated using Accutase and plated on a Geltrex-coated well at a density of 40,000–68,000 cells/cm^2^ (depending on the cell line) in Stemflex with 4 μM ROCK inhibitor Y-27632 (10005583; CAYMAN CHEMICAL, MI, USA). On day 1, we added mesoderm induction medium consisting of N2B27 medium (1:1 mixture of DMEM:F12(1:1) with Glutamax (10565018; Thermo Fisher Scientific) and Neurobasal medium (21103049; Thermo Fisher Scientific, MA, USA) supplemented with N2 (17502048; Thermo Fisher Scientific) and B27 (17504044; Thermo Fisher Scientific, MA, USA)), 8 μM CHIR99021 (C2943; TCI, Japan), and 25 ng/mL bone morphogenetic protein 4 (BMP4) (314-BP; R&D Systems, MN, USA). On days 4 and 5, the mesoderm induction medium was replaced with VSMC induction medium consisting of N2B27 medium with 12.5 ng/mL platelet-derived growth factor-BB (100-14B; PeproTech, NJ, USA) and 12.5 ng/mL activin A (338-AC; R&D systems). On day 6, differentiated VSMCs were dissociated and plated on type I collagen (354265; Corning, NY, USA)-coated plates in Medium 231 (M231500; Thermo Fisher Scientific) with 1% smooth muscle differentiation supplement (S0085; Thermo Fisher Scientific).

For differentiation to ECs, iPSCs were dissociated using Accutase and plated on a Geltrex-coated well at a density of 25,000–50,000 cells/cm^2^ (depending on the cell line) in Stemflex with 4 μM ROCK inhibitor Y-27632. On day 1, the medium was replaced with DMEM/F12 Glutamax medium supplemented with B27 minus Vitamin A (12587010; Thermo Fisher Scientific), 1 μM CHIR99021, 10 nM retinoic acid (RA) (R2625; Sigma-Aldrich, MO, USA), 1 ng/mL BMP4, and 100 ng/mL basic fibroblast growth factor 2 (bFGF2) (19155-36; Nacalai, Japan). After 24 h, the medium was replaced with DMEM/F12 Glutamax medium supplemented with B27 minus Vitamin A, 3 μM CHIR99021, 100 ng/mL bFGF, and 25 ng/mL BMP4 and cultured for 3 days to induce lateral plate mesoderm. Then, the cells were dissociated with Accutase and plated on Laminin 411 (892042; Nippi, Japan)-coated plates in StemPro-34 SFM (10639011; Thermo Fisher Scientific) supplemented with 4 μM ROCK inhibitor Y-27632, 50 ng/mL recombinant human vascular endothelial growth factor (QK048-0025; MBL, Japan), 10 ng/mL bFGF2, 10 μM DAPT (552-29251; Wako, Japan), 20 ng/mL BMP4, and 10 μM IWR-1 (I0161; Sigma-Aldrich). The cells were cultured for 4 days to complete the differentiation. On day 9, the differentiated ECs were dissociated and plated on fibronectin (356008; Corning, NY, USA)-coated plates in endothelial growth medium (CC3162; Lonza, Switzerland).

All cells were cultured at 37°C in a humid atmosphere with 5% CO_2_. The experiments described in this manuscript were performed with iPSC-VSMCs and ECs in passages 3 and 4. The antibiotics Zellshield (13-0150; Funakoshi, Japan) and penicillin–streptomycin (09367-34; Nacalai, Japan) were added to Stemflex and all other media, respectively.

### Preparation of collagen solutions

Two concentrations of collagen pre-gel solution were prepared for the vascular tube fabrication. All solutions were mixed on ice to prevent the unexpected solidification of collagen. Then, 10X Hanks’ balanced salt solution (H4641; Sigma-Aldrich), 1 M HEPES solution (H0887; Sigma-Aldrich), 1 M NaHCO_3_ solution (144-44-8; Nacalai), distilled water, and 5 mg/mL acid solubilized collagen solution (IAC-50; KOKEN, Japan) were mixed to reconstitute the collagen pre-gel solution. These five solutions were mixed at volume ratios of 10:1:1:68:20 and 10:1:1:8:80 to prepare 1 and 4 mg/mL collagen solutions, respectively, which were used for VSMC fiber formation and tube fabrication, respectively.

### Fabrication of the ViTAP model

A polydimethylsiloxane (SILPOT 184; Dow TORAY, Japan) mold with a thin groove was fabricated using a 3D-printed PLA mold, and tungsten wires (φ0.2 mm and φ0.5 mm, 461267 and 462384; Nilaco, Japan) were placed on the mold. Then, VSMCs differentiated from iPSCs were dissociated from the dishes by trypsinization with trypsin EDTA (32777-44; Nacalai) and suspended in 1 mg/mL collagen pre-gel solution. The cell-suspended solution was soaked into the groove of the mold. The mold with cells was cultured for 24 h at 37℃ to form a high-density fiber. The contracted VSMC fiber was then rolled up by tweezers to form a helical shape. Connectors made of glass capillaries (83-0528; Sansho, Japan) and silicone tubes (285-01781; Hagitec, Japan) were inserted at both ends and embedded within a 4 mg/mL collagen pre-gel solution in a microfabricated PLA mold. After incubating for an hour at 37℃ to solidify collagen, the PLA mold and tungsten wires were removed to obtain the tube-shaped model. Finally, ECs differentiated from iPSCs were dissociated from the dishes by trypsinization with trypsin EDTA and seeded into the inner wall using a micro syringe (1725TLL; Hamilton, NV, USA) and an injection needle (NN-2516R; Terumo, Japan) to build the vascular model.

### Fluorescent staining

The actin filaments of VSMCs were stained with a live-cell actin probe (SPY555-actin; Cytoskeleton, CO, USA). The probe was directly added to the culture medium including vascular models at 1:1000 dilution ratio. After incubating for an hour, the probe was removed by changing the medium to finish the staining.

For immunostaining, the cells were immersed in a 4% paraformaldehyde solution for 15 min at 37℃ for fixation. Then, the fixed cells were incubated in PBS supplemented with 0.1% Triton X-100 (PBS-T) (X-100; Sigma-Aldrich) including 5% donkey serum (D9663; Sigma-Aldrich) for blocking and permeabilization for an hour at 22–24℃. Subsequently, the cells were reacted with primary antibodies in 5% donkey serum in PBS-T overnight at 4℃. The following day, the cells were washed with PBS and then incubated with secondary antibodies in 5% donkey serum in PBS-T for an hour at room temperature in the dark. After the incubation, the buffer was removed and replaced with PBS supplemented with Hoechst 33342 (H3570; Thermo Fisher Scientific) at room temperature for 10 min in the dark. The cells were then washed with PBS and mounted with an aqueous-based mounting medium to prevent fading. The antibodies used are listed in the key resources table.

### Image analysis

The alignment of actin filaments in the VSMCs was analyzed using Fiji, a distribution of ImageJ (National Institutes of Health, MD, USA), with LPixel and directionality plugins. First, the fluorescently stained actin filaments were captured in detail by a confocal microscope (C2; Nikon, Japan). Then, the image was binarized by the image thresholding of ImageJ. Subsequently, LPixel was used to extract the morphologies of actin filaments as skeleton images. This step was performed in line extract mode by using a multidirectional non-maximum suppression algorithm. Finally, the direction of extracted lines was analyzed with the directionality plugin to evaluate the alignment of cells.

### Set up and experimental flow for hormonal vasoconstriction and vasodilation

The ViTAP models were placed in a stage-top incubator (STRG-KIW; Tokaihit, Japan) in a microscope (BZ-X810; KEYENCE, Japan) to observe the vascular contraction. Before the experiment, the tube was acclimated to the chamber for an hour. When the time-lapse imaging was started, the vasoconstrictors and vasodilators (U46619 (ab144540; Abcam, MA, USA), endothelin-1 (ab120471; Abcam), angiotensin II (336-40011; Wako, Japan), adrenaline (E4642; Sigma-Aldrich), noradrenaline (A7256; Sigma-Aldrich), and prostaglandin I₂ (P6188; Sigma-Aldrich) were exposed at the desired concentration to vascular models. Then, 100 mM KCl was added to measure the maximum contraction ratio of each model for normalization. The relative contraction was calculated by dividing each contraction ratio by the maximum contraction ratio.

The pretreatment of ViTAP models with nifedipine (N7634; Sigma-Aldrich) or MA-5 was started 30 min before the experiments.

### Set up and experimental flow for pressure-derived vasodilation

A pressure-based pump with a microfluidic flow control unit (Flow EZ; Fluigent, France) was used to apply 50 mmHg of physiological blood pressure to the vascular model. The ViTAP models were placed in an originally microfabricated perfusion dish, which has silicone connectors to an external pump through the wall, and set on a live-cell imaging microscope (CytoSMART Lux3 FL; Shoshin EM, Japan). Then, the connectors of the dish were attached to the pressure pump through silicone tubes.

The expansion behavior was analyzed using motion tracking. First, the vascular model was monitored and recorded by a PC connected to the microscope while applying 50 mmHg of pulsatile pressure. Next, arbitrary tracking points on both ends of the outer border of VSMC layers were chosen on the captured video, and the position was traced using video analysis software (VW-9000 Motion Analyzer; KEYENCE). Finally, the change in the diameter was calculated to see the deformation behavior. If vasoconstrictors were simultaneously exposed to the vascular model, the chemicals were directly added to the vascular model on the perfusion dish.

### RNA isolation, cDNA synthesis, and qPCR

RNA extraction was conducted using Sepasol reagent (09379-97; Nacalai) in accordance with the manufacturer’s instructions. cDNA synthesis was performed using ReverTra Ace qPCR RT master mix in accordance with the manufacturer’s instructions (FSQ-301; TOYOBO, Japan). WRN, p16, and b-actin qPCR were performed using the primer sets listed in the key resources table in conjunction with SYBR Green qPCR mix on a QuantStudio3 Real-Time PCR System (Thermo Fisher Scientific). Expression data are presented after calculating the relative expression compared with the endogenous control gene b-actin.

### Spatial transcriptome analysis

The vascular model was fixed in 10% neutral buffered formalin and embedded with paraffin through general dehydration, clearing, and paraffin infiltration using ethanol and xylene to create a formalin-fixed paraffin-embedded (FFPE) tissue block. Then, the FFPE tissue block was cooled and cut into 3.5–5 µm sections by using a microtome.

The sections were placed on Superfrost Plus Slides (Epredia, 6776214) in accordance with the Tissue Preparation Guide (CG000518 Rev B, 10X Genomics, CA, USA) and adhered by 3 h of warming at 42℃ and desiccating overnight. The sections were then deparaffinized, H&E stained, imaged, and decrosslinked in accordance with the Visium CytAssist Spatial Gene Expression for FFPE User Guide (CG000520 Rev B, 10X Genomics). Brightfield histology images were taken using a microscope (BZ-X700; Keyence, Japan). Raw images were stitched using BZ-X analyzer software (Keyence, Japan) and exported as TIFF files. Then, the samples underwent probe hybridization, ligation, and release and extension to generate spatially barcoded and ligated probe products, which can be carried forward for library preparation (CG000495 Rev C, 10X Genomics). After pre-amplification followed by cleanup by SPRIselect, qPCR was performed to determine the Sample Index PCR cycle number for gene expression libraries. Finally, sample index PCR was conducted to generate library molecules followed by cleanup with SPRIselect and quality control of library construction with a bioanalyzer (CG000495 Rev C).

Libraries were sequenced on a NovaSeq 6000 System (Illumina, CA, USA) using a NovaSeq S4 Reagent Kit (200 cycles, 20027466; Illumina) at a sequencing depth of approximately 250–400 M read pairs per sample. Sequencing was performed using the following read protocol: read 1, 28 cycles; i7 index read, 10 cycles; i5 index read, 10 cycles; read 2S, 50 cycles. Raw FASTQ files and histology images were processed by sample with the Space Ranger software (ver 3.0.1, 10X Genomics) against the Cell Ranger mm10 reference genome “refdata-gex-GRch38-2020-A,” available at: https://cf.10xgenomics.com/supp/spatial-exp/refdata-gex-GRch38-2020-A.tar.gz. Seurat v4 was used for downstream analysis. The datasets were individually subjected to FindIntegrationAnchors and IntegratedData functions and merged. The batch effect of the merged data was corrected by harmony. The merged data was used for dimensionality reduction and cluster detection. Differentially expressed genes were detected using FindMarkers in Seurat (log2fc.threshold > 0.20, p_val < 0.05).

The differentially expressed genes were input into Enrichr to perform gene enrichment analysis and highlight signaling pathways^47–49^. The adjusted p-value was computed using the Benjamini–Hochberg method for correction for multiple hypotheses testing.

### Metabolomic analysis

Five vascular tubes were pooled as one sample, estimated to contain 1 × 10⁶ cells, to ensure a sufficient number of cells for metabolomic analysis. The untargeted metabolic profiling platform was conducted by Metabolon. The data were normalized based on the protein level of each sample. As two different models (WT and Werner) were aligned as biological replicates, Welch’s two-sample t-test was employed to comprehensively identify metabolites that differed significantly between experimental groups.

### Statistical analysis

Prism 7 (GraphPad, MA, USA) was used to create graphs and perform statistical analyses. Statistical analysis was performed using unpaired, two-tailed Student’s t-test, one-way ANOVA followed by Tukey’s multiple comparison test, or two-way ANOVA followed by Sidak’s multiple comparison test. Non-normally distributed data were analyzed using the Mann–Whitney test. For all bar graphs, data are represented as mean ± S.D. Statistical significance was considered at *p* < 0.05.

### Data and code availability

The spatial transcriptomic data have been deposited in GSE302978 (https://www.ncbi.nlm.nih.gov/geo/query/acc.cgi?acc=GSE302978).

## ACKNOWLEDGMENTS

This work was partly supported by JSPS KAKENHI (Grant Nos. JP23K17184, JP22J01133, and JP24K11379) and TERUMO LIFE SCIENCE FOUNDATION, AMED (Grant No. 24ek0210168h0003 and 24fk0108655h0003). This work was also partly supported by Biomedical Research Unit of Tohoku University Hospital. The authors would like to thank Misuzu Naka and Keiko Suto (Tohoku University) for their assistance in the culture and differentiation of iPSCs and Kiyomi Kisu (Tohoku University) for her guidance and lecture on FFPE tissue section preparation for spatial transcriptomic analysis. The authors also would like to thank Ben Pope (MelliCell, Inc) for his detailed revision on our manuscript. We also acknowledge the depositor of the WS2-C5 cell line, Dr. Miyoshi Hiroyuki, and the establisher, Dr. Atsushi Miyawaki, for their huge contribution to aging research.

## AUTHOR CONTRIBUTIONS

S.I., H.O., and T.T. conceived the project and designed the study. S.I., R.U., and M.K. performed most of the experiments and analyzed data. J.M. conducted the motion capture to analyze the deformation behavior. M.K. and S.N. designed and performed the spatial transcriptomic analysis. T.K. contributed to the design of the methodology, especially in the differentiation of iPSCs. T.T., H.O., and T.A. provided general feedback on the results and contributed to the leadership of the research. S.I., T.T., and T.A. acquired the funding. S.I. and T.T. wrote the paper. All authors reviewed and commented on the paper.

## DECLARATION OF INTERESTS

The authors declare no competing interests.

## SUPPLEMENTAL INFORMATION

Document S1. Figures S1–S5

Video S1. The vasoconstriction behavior of the ViTAP model after exposure to 200 nM of U46619, a thromboxane A2 receptor agonist. (×300 speed), related to Figure 2

Video S2. The vasodilation behavior of the ViTAP model after exposure to 1 μM of prostacyclin (prostaglandin I₂). (×300 speed), related to Figure 2

Video S3. The vasodilation behavior of the ViTAP model under 50 mmHg of pulsatile pressure. (×1 speed), related to Figure 2

## Notes

### Competing Interest Statement

The authors have declared no competing interest.

