## Supplemental file, movies for "Bioengineered iPSC Vessels Recapitulate Human Vascular Physiological Function and Aging Phenotypes": Supplemental information_CSC_ShunItai.pdf

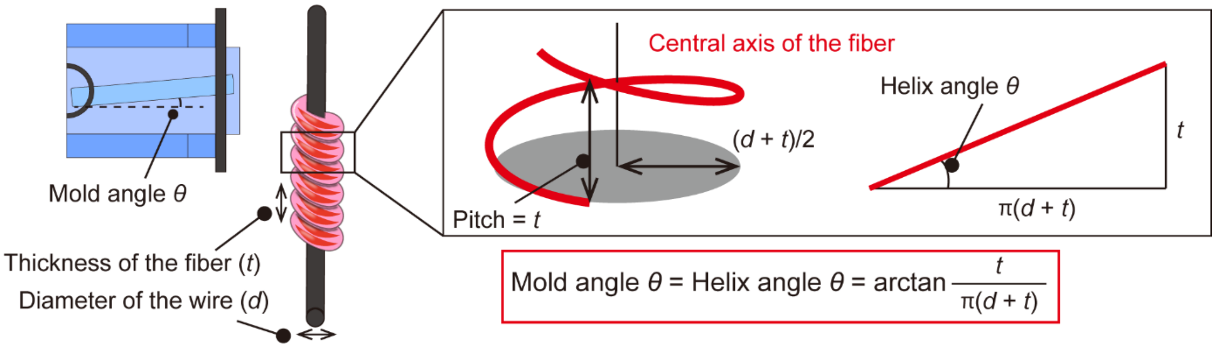


**Figure S1**. Design of the microfabricated mold. The mold angle was calculated based on the fiber thickness and wire diameter to achieve a uniform roll-up of the VSMC fiber into a helical form.

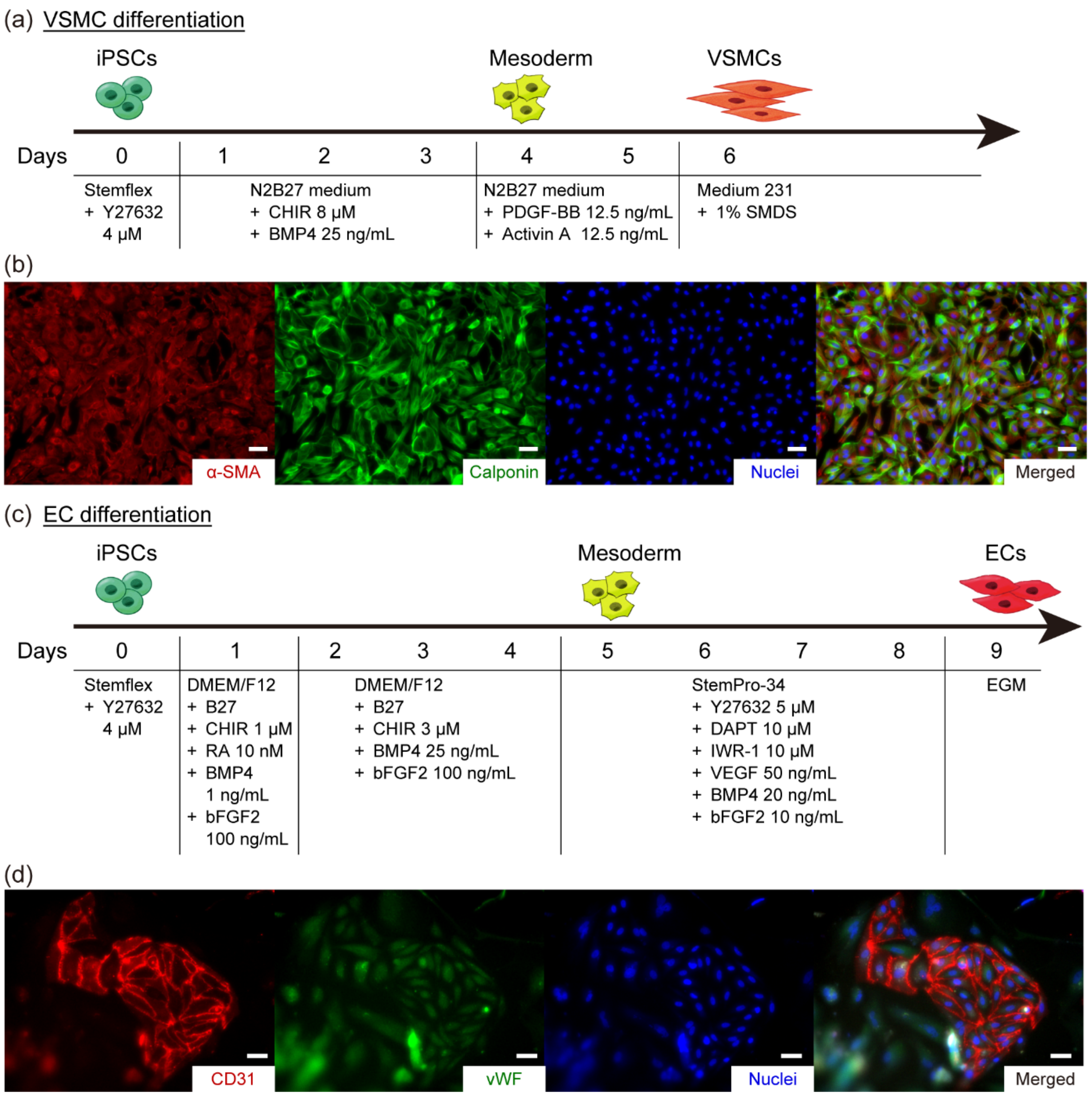


**Figure S2**. Differentiation of iPSCs (a) Protocol to induce VSMCs. (b) Immunostaining of differentiated VSMCs. (c) Protocol to induce ECs. (d) Immunostaining of differentiated ECs (Scale bars: 50 μm).

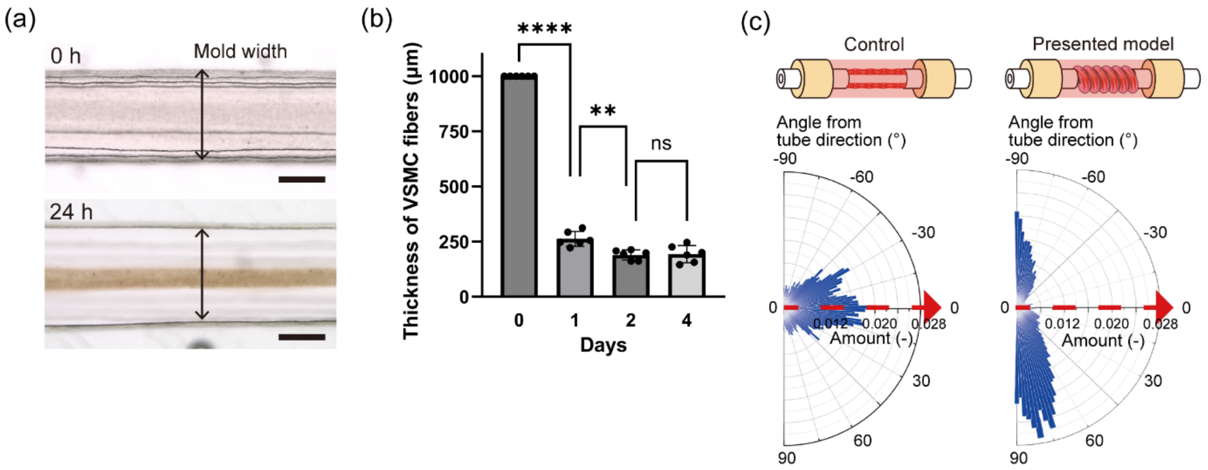


**Figure S3**. Supplemental fabrication results of the ViTAP model. (a) Microscopic images of VSMC fiber formation (Scale bars: 500 μm). (b) Change in diameter of VSMC fibers ($n$ = 6). Cells formed high-density fibers within 24 h of incubation. (c) Alignment analysis of actin filaments. Compared with the control model, our model showed the radial alignment of VSMCs (** $p < 0.01$, **** $p < 0.0001$).

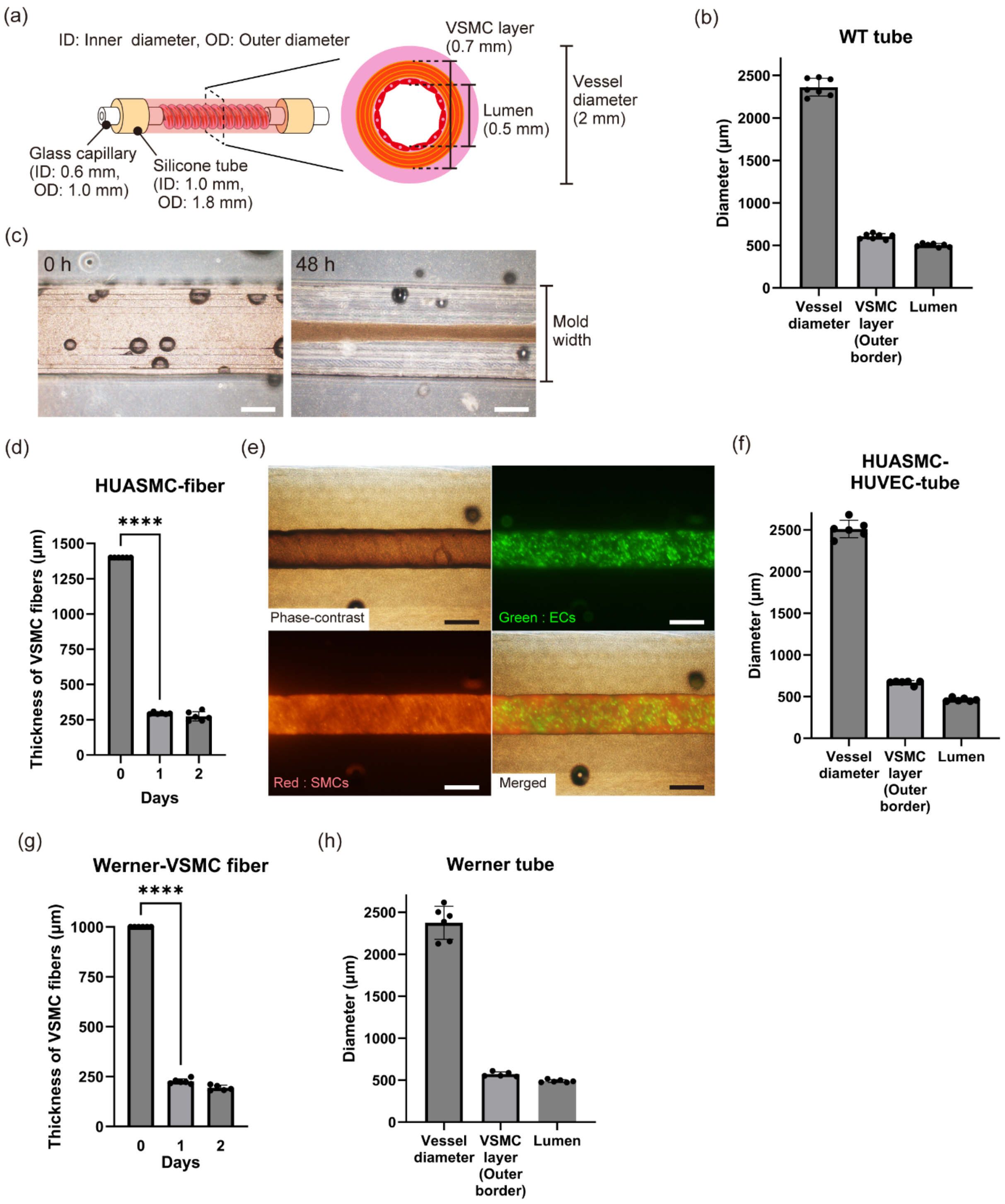


**Figure S4**. (Continued on next page)

**Figure S4**. Supplemental fabrication results of the Vascular Induced Pluripotent Stem Cell (iPSC) Tube Achieving Physiological Deformation (ViTAP) model with various cell sources. (a) Design of the size of the vascular model used for the experiments in this paper. (b) Diameters of the fabricated ViTAP model ($n$ = 6). The coefficient variations were under 5% for all diameters. (c) Fiber formation results using human umbilical artery smooth muscle cells (HUASMCs) (Scale bars: 500 μm). (d) Fiber formation results using HUASMCs ($n$ = 6). (e) Fabricated tissue using HUASMCs and human umbilical vein endothelial cells (HUVECs). Co-culture of circumferentially aligned SMCs and ECs was achieved (Scale bars: 500 μm). (f) Diameter distribution of the HUASMC-HUVEC model ($n$ = 6). (g) Fiber formation results of the Werner model ($n$ = 6). (h) Diameter distribution of the Werner model ($n$ = 6) (**** $p < 0.0001$).

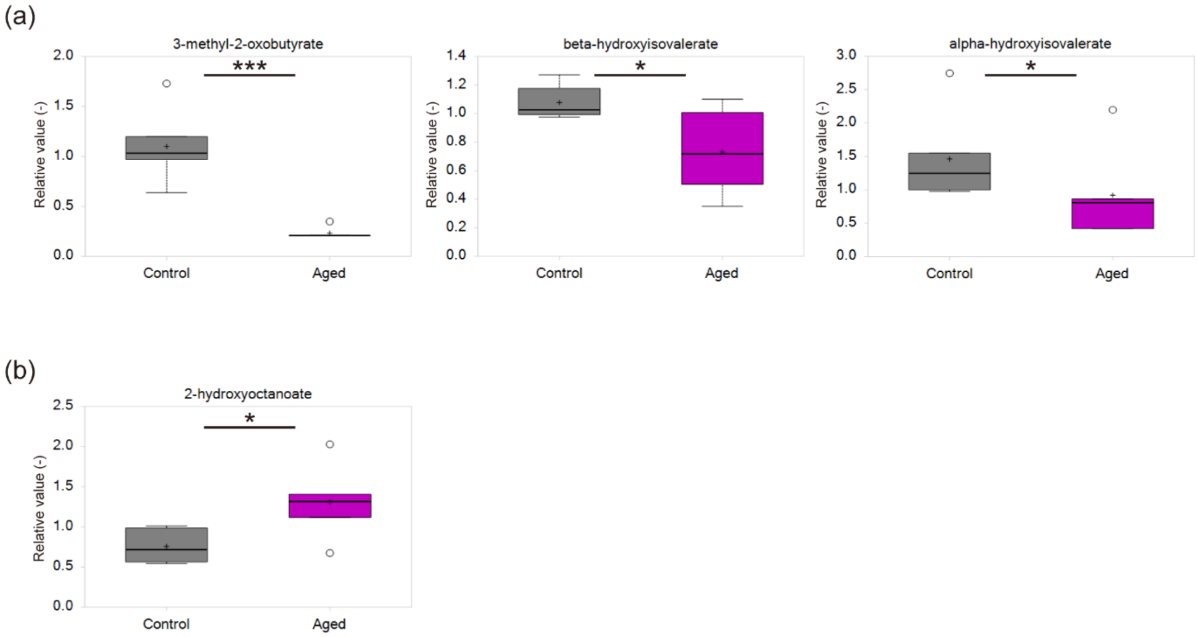


**Figure S5**. Supplemental results of the comprehensive analysis of vascular aging on the ViTAP model ($n$ = 6). (a) Metabolite abundance of the group of branched-chain α-ketoacids. (b) Metabolite abundance of the group of monohydroxy fatty acids (* $p < 0.05$, *** $p < 0.001$).
